# Temporal and spatiotemporal perturbations in paced finger tapping point to a common mechanism for the processing of time errors

**DOI:** 10.1101/690834

**Authors:** Sabrina L. López, Rodrigo Laje

## Abstract

Paced finger tapping is a sensorimotor synchronization task where a subject is instructed to keep pace with an external metronome, as when following along with the beat of music, and the time differences (asynchronies) between each stimulus and its response are recorded. The usual way to study the underlying error correction mechanism is to make unexpected temporal perturbations to the stimuli sequence and then let the subject recover average synchronization. A critical but overlooked issue in traditional temporal perturbations, however, is that at the moment of perturbation two things change: both the stimuli period (a parameter) and the asynchrony (a variable). In terms of experimental manipulation, it would be desirable to have separate, independent control of parameter and variable values. In this work we perform paced finger tapping experiments combining simple temporal perturbations (tempo step change) and spatial perturbations with temporal effect (raised or lowered point of contact). In this way we decouple the parameter-and-variable confounding of traditional temporal perturbations and perform novel perturbations where either the parameter only changes or the variable only changes. Our results show nonlinear features like asymmetry and are compatible with the idea of a common mechanism for the correction of all types of asynchronies. We suggest taking this confounding into account when analyzing perturbations of any kind in finger tapping tasks but also in other areas of sensorimotor synchronization, like music performance experiments and paced walking in gait coordination studies.

## 1. Introduction

Sensorimotor synchronization (SMS), the mainly specifically human ability to keep synchrony with an external periodic metronome, is the basis of all music and dance [ReppSu2013, Patel2009, Schachner2009, Zatorre2007]. It is a spontaneous behavior found in humans and, despite its simplicity, the mechanisms underlying it remain mostly unknown; the processing of temporal information is an open area of research in neuroscience and how time is represented and manipulated in the brain is still one of the most elusive concepts, particularly in the millisecond timing range (hundreds of milliseconds) [IvrySpencer2004, IvrySchlerf2008, BuonomanoLaje2010, Grondin2010, Paton2018].

The most used experimental paradigm in SMS is auditorily paced finger tapping, a task where a subject is instructed to move a finger (tap) in synchrony with a periodic stimulus sequence of short tones (beep) as in keeping pace with music (Figure fig.tappinga). The main observable, both for behavior description [Chen1997] and for mathematical modeling [ReppSu2013], is the asynchrony *e*_*n*_ = *R*_*n*_ − *U*_*n*_, that is the difference between the occurrence time of each response R_n_ and the occurrence time of its corresponding stimulus *U*_*n*_. It is simple to demonstrate the existence of an underlying error correction mechanism---if the stimuli sequence is suddenly silenced while the subject keeps tapping, the “virtual asynchronies” between responses and extrapolated beeps become very large very soon drifting away from average synchrony. In order to understand the inner workings of such error correction mechanism, temporal perturbations to the stimuli sequence are usually performed and then study the resynchronization [Bavassi2013]. Traditional temporal perturbations consist in for example an abrupt change in the period of the sequence from a given beep on, which is known as a tempo step change (Figure fig.tappingb).

**Figure fig.tapping.**
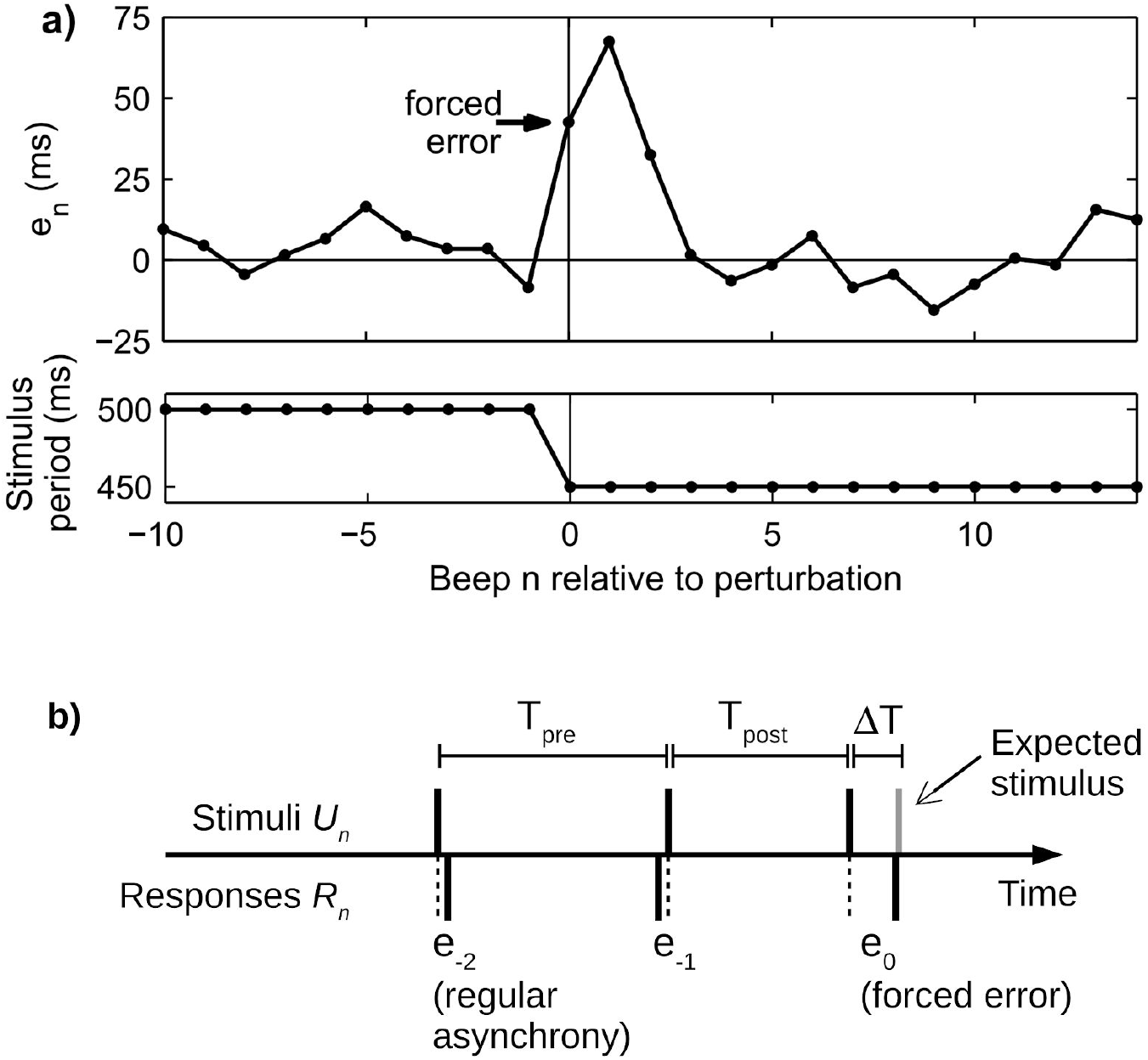
Paced finger tapping task with an unexpected tempo perturbation. (a) Asynchrony time series (top) from a single trial from one subject showing the forced error produced by a tempo decrease (bottom) occurring at n=0. (b) Schematic and definitions. As the perturbation DeltaT is unexpected, a change in the variable e_0 occurs in addition to the change in the parameter T.

There is, however, an overlooked issue with traditional perturbations. Let’s consider a simple perturbation like the mentioned tempo step change. At step n when the perturbation occurs, both the parameter value (stimuli period) and the variable value (asynchrony) change, because the stimulus occurrence time *U*_*n*_ changes and thus *e*_*n*_ changes instantaneously and arbitrarily---importantly, without involving any dynamics of the underlying system (Figure fig.tappingb). The critical difference between period and asynchrony is that the former has no dynamics by itself and its value is up to the experimenter, while the latter is subject to the dynamics of the system under study; they might have, for instance, different neural correlates. Note that this issue appears in any traditional perturbation that modifies any stimulus occurrence in the sequence, like for instance the above-named tempo step changes [ReppKeller2004, Thaut1998], but also sigmoidal changes [SchulzeCordesVorberg2005, PecenkaKeller2009, vanderSteen2015], linear changes [Loehr2011], quadratic changes [Pecenka2013], sinusoidal changes [Michon1967, PecenkaKeller2011], random changes [Michon1967, HaryMoore1987, MadisonMerker2004, ReppPenel2004], phase shifts [Repp2001phaseshifts], event onset shifts [Repp2002phasecorrection, Repp2002automaticity, Flach2005], and adaptively timed sequences [ReppKeller2008, Repp2012, Mills2015]. Up to our knowledge, this is the first work to note this issue.

Beyond the finger tapping literature, research in other areas within SMS are quite likely impacted by this parameter-variable confounding. Two areas are easily spotted. First, research on music performance and perception also relies on perturbation experiments and ecological experimental conditions with natural variability in the stimuli, like accelerando, ritardando, and natural expression [Rankin2009], groove and microtiming [Madison2011], musicians following a conductor [Luck2006], phase shift and tempo perturbations [Levitin2018, Zanto2005, LargeKelso2002], auditory delayed feedback [Pfordresher2006], and even on theoretical works about oscillator models of synchronization to a musical beat [LargePalmer2002]. Second, research on gait coordination also makes use of experimental designs that might be confounding parameter and variable perturbations, like phase shift perturbations [Roerdink2009, Pelton2010], interpersonal coordination during side-by-side walking [vanUlzen2010], and varying belt speeds and directions in treadmill walking [Choi2007].

An analogy may help convey our point and our proposal. Let’s consider the circadian clock, a well-characterized genetic oscillator that in mammals is in the suprachiasmatic nucleus of the hypothalamus. The circadian clock displays an autonomous oscillatory activity with a period of around 24 hs capable of synchronizing to the daily cycle mainly through the light/dark stimulus [Golombek2010, Laje2018]. In a minimalistic description, the system works as follows: a gene produces a protein (BMAL/CLOCK) that activates a second gene, which in turn produces a second protein (PER/CRY) that inactivates the first gene. In this system the period of the external light/dark stimulus is a parameter, while the concentrations of CLOCK/BMAL and PER/CRY proteins are the variables whose time evolution is set by the dynamics of the system. Both the parameter and the variables may be experimentally manipulated independently: the parameter value can be modified by changing the period of the external light/dark stimulus (called T-cycle [Golombek 2013, Plano2010]), and the value of the variables can be modified for instance by applying an acute dose of inhibitor PF-670462 that modifies the concentration of PER [Kim2013].

It would be desirable in paced finger tapping to manipulate parameter and variable values independently. In this work we perform paced finger tapping experiments with simple temporal perturbations (traditional tempo step changes), and also novel spatial perturbations with temporal effect (raised or lowered point of contact) and combined perturbations with the aim of probing the error correction mechanism. This allows us to decouple the effect of traditional temporal perturbations and perform novel perturbations where the parameter only changes (a change in stimuli period without a change in asynchrony) and where the variable only changes (a change in asynchrony without a change in period). Our results show nonlinear effects even when the perturbations are 10% of the stimulus period and are compatible with the idea that the origin of the asynchrony doesn’t influence the subsequent resynchronization. The issue of whether motor timing and sensory timing depend on the same neural circuits is still open [BuonomanoLaje2010], but our results suggest that asynchronies produced by purely temporal perturbations and those produced by spatiotemporal perturbations are processed by a common error correction mechanism.

## 2. Results

We performed an auditorily paced finger tapping experiment with perturbations, where the subject is instructed to keep in synchrony at his/her best and keep tapping to resynchronize in case a perturbation appears at a random beep. Perturbations can be any of the following: a) simple temporal perturbations “T” (traditional tempo step change perturbations by an amount +/−DeltaT); b) simple spatial perturbations “S” that have temporal effect, achieved by raising (+) or lowering (−) the point of contact where the subject has to tap and thus advancing or delaying the time of contact; c) combined simultaneous perturbations “ST”. Perturbations were classified according to two criteria: by size (small and large perturbations), and by type (simple temporal +/−T; simple spatial +/−S; combined analogous +S+T and −S−T; and combined opposite +S−T and −S+T). Note that in the simple spatial +/−S perturbations the stimuli period doesn’t change and thus they are variable-only perturbations; in the combined opposite +S−T and −S+T the stimuli period changes but the asynchrony doesn’t because both components compensate each other on average at the perturbation beep, and thus they are parameter-only perturbations. See Methods for a detailed description and rationale.

Figure fig.todas shows the average time series for every condition. At the perturbation beep n=0 a forced error occurs as the perturbations are unexpected. Small perturbations take more steps to recover than large perturbations.

**Figure fig.todas.**
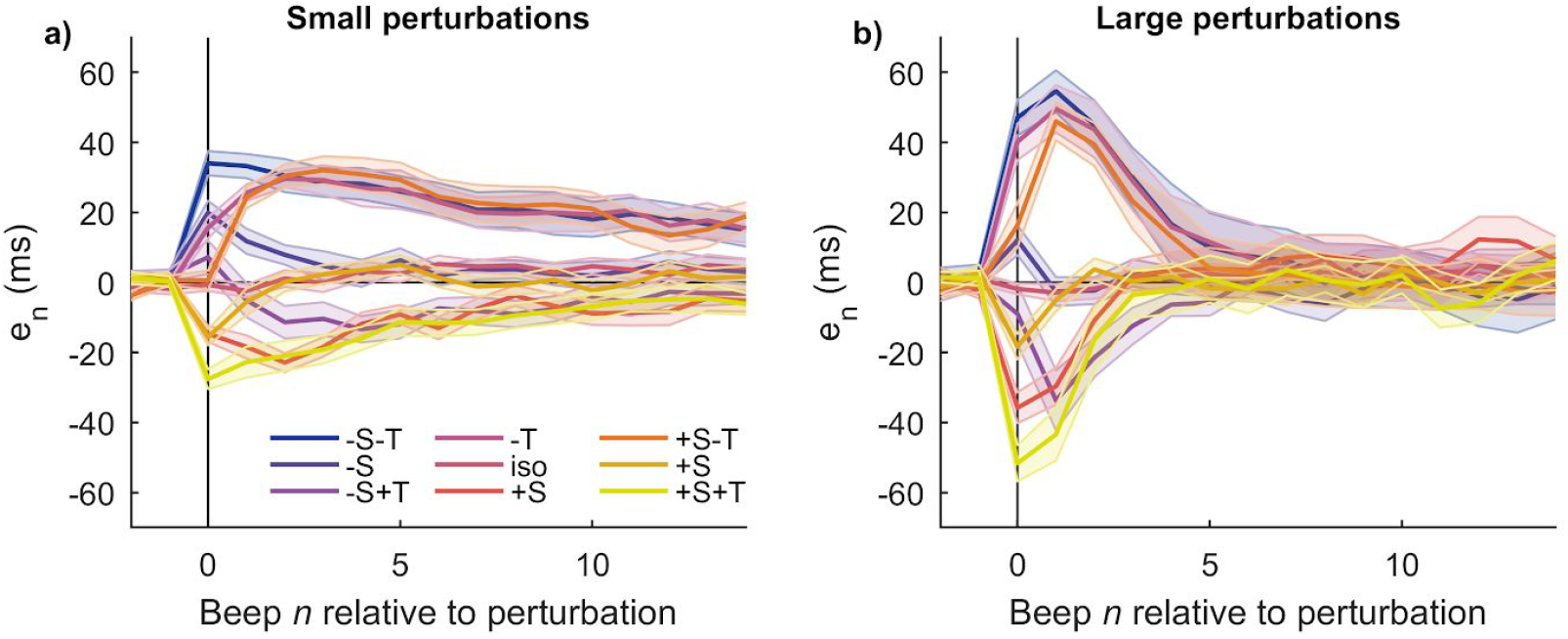
Resynchronization time series for every condition averaged across subjects. Perturbation occurs at n=0. Single trials were vertically shifted before averaging such that its pre-perturbation baseline is zero. (a) Small perturbations. (b) Large perturbations. All traces are mean +/− standard error across subjects.

One of the effects of a tempo step change perturbation in paced finger tapping is a shift in baseline relative to its pre-perturbation value [Bavassi2013]. Such baseline, which represents the state of the system in stationary conditions, is the extensively studied “negative mean asynchrony” or NMA [Repp2005]. In this work, on the contrary, we are interested in analyzing the system dynamics right after being perturbed and how it converges to its post-perturbation value [Bavassi2013]. In order to do that, from here on we choose to plot every time series after subtracting its post-perturbation baseline such that all time series converge to zero after perturbation (see Methods).

### 2.1. The response to simple perturbations is asymmetric

In Figure fig.simplesa and b we show the averaged time series for perturbations +T and −T. Large perturbations produce asymmetric responses--the response to the positive perturbation converges to zero more rapidly than the negative one, in accordance with previous reports for this type of perturbation [Thaut1998, Bavassi2013]. Small perturbations, on the other hand, do not show asymmetry. Both responses are consistent with a nonlinear underlying system, where nonlinear behavior like asymmetry is evident only when the perturbation size is large enough [Loehr2011].

**Figure fig.simples.**
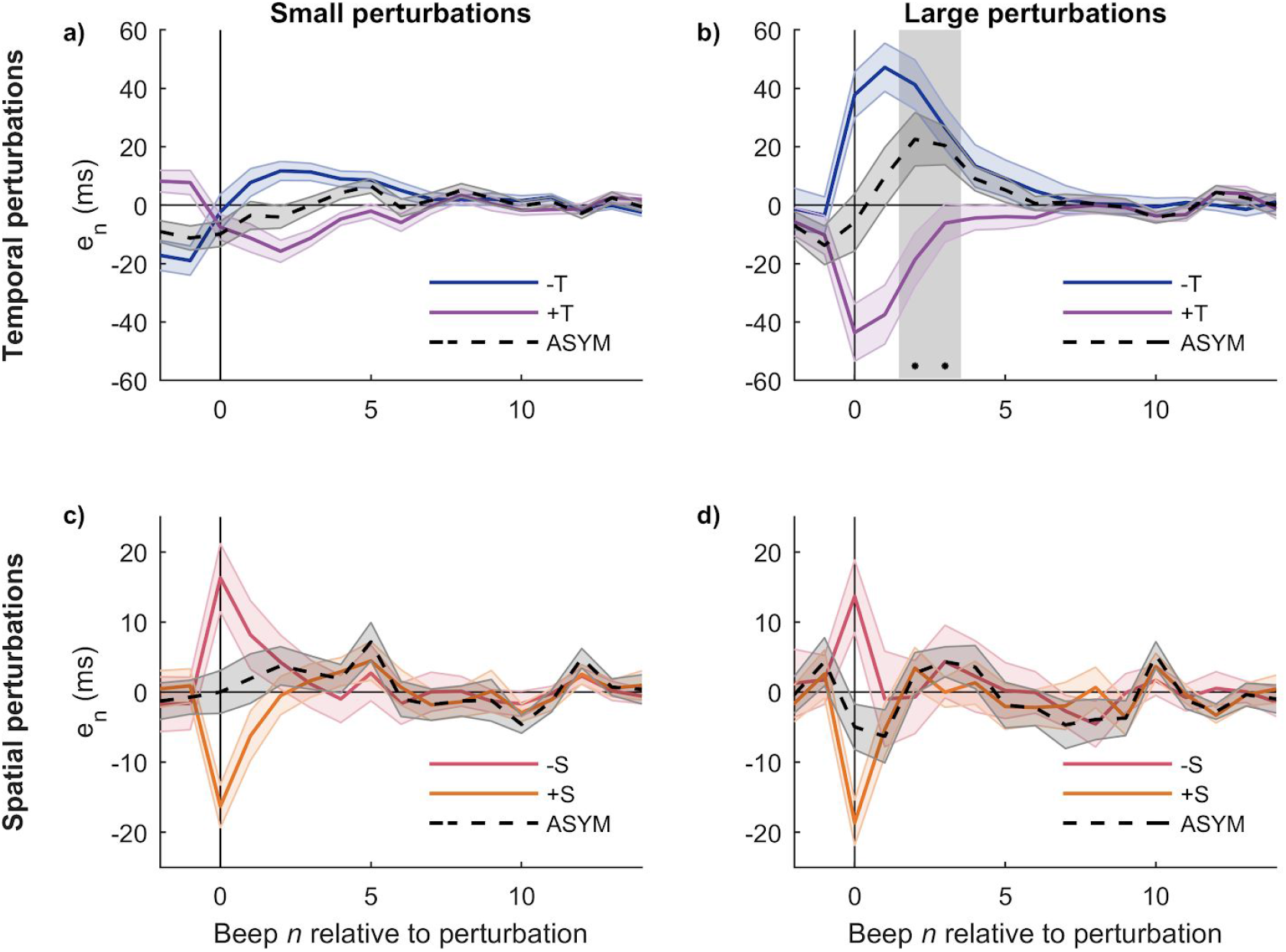
The response to large temporal perturbations is asymmetric. (a) and (b) Response to traditional tempo step change perturbations (+/−T) and degree of asymmetry between them (ASYM). (c) and (d Response to novel spatial perturbations (+/−S) and degree of asymmetry between them (ASYM). Left: small perturbations; Right: large perturbations. Large temporal perturbations produce asymmetric responses, and the grey area indicates steps where the asymmetry is significative (p=0.0025 each). All traces are mean +/− standard error across subjects.

Spatial perturbations +S and −S, on the contrary, produce responses that are mostly symmetric (Figure fig.simplesc and d). It is worth noting that the asynchrony values produced by +/−S perturbations at the perturbation beep are relatively small and similar in size to those produced by the small +/−T perturbations, which also have symmetric responses (Figure fig.simplesa). Spatial perturbations +/−S are novel in that they produce a change in the value of the variable (asynchrony *e*_*n*_) without changing the stimulus period.

### 2.2. The response to combined perturbations is asymmetric

In this section we analyze the degree of asymmetry of the combined perturbations, and for that we group them in “analogous” (+S+T and −S−T, because their components S and T individually would produce asynchronies of the same sign) and “opposite” (+S-T and −S+T, whose components would individually produce asynchronies of opposite signs; see Methods). Figure fig.combinadasa and b shows that the large analogous perturbations produce asymmetric responses; when compared to the large temporal perturbations (+T y−T, Figure fig.simples) the degree of asymmetry seems greater in the analogous perturbations (−S−T and +S+T) and lesser in the opposite perturbations (−S+T and +S−T).

**Figure fig.combinadas.**
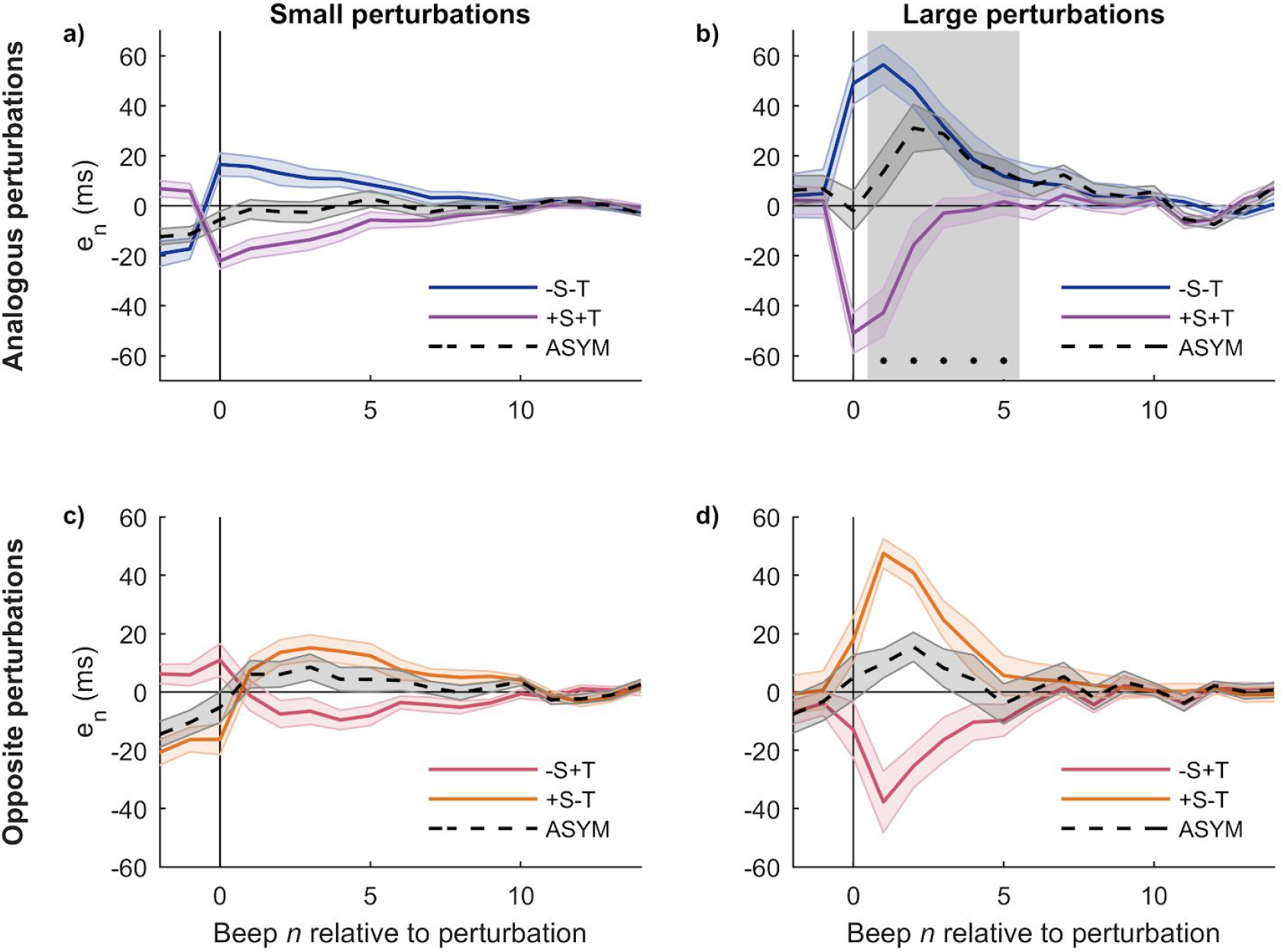
Large combined perturbations produce asymmetric responses. (a) and (b) Response to analogous perturbations −S−T and +S+T (whose components S and T would individually produce asynchronies of the same sign) and degree of asymmetry (ASYM). (b) and (c) Response to opposite −S+T and +S−T (whose components S and T would individually produce asynchronies of opposite signs) and degree of asymmetry (ASYM). Left: small perturbations; Right: large perturbations. When compared to the large temporal perturbations of the previous section, analogous perturbations produce more asymmetric responses while opposite perturbations produce less asymmetric responses. Mean +/− standard error across subjects. The grey area indicates steps when the asymmetry is statistically significant (p=0.022, 0.0025, 0.0025, 0.0067, 0.022, respectively).

We note that the analogous perturbations produce asynchronies with values at the perturbation beep (n=0) greater than the temporal perturbations, while the opposite perturbations produce values lesser than the temporal perturbations (Wilcoxon sign-rank test, significant after Bonferroni correction; simple temporal vs. analogous p=0.0134; simple temporal vs. opposite p=0.00073; analogous vs. opposite p=0.00048). The result in this section may be understood within the same framework that the result from Section 3.1: it is the behavior of a nonlinear underlying system whose asymmetric response is only evident when the asynchrony values are large enough, and whose degree of asymmetry is larger for greater asynchronies.

### 2.3. Simple perturbations are additive

One of the aims for proposing the combined perturbations (+/−)S(+/−)T was to compare every combination with the simple perturbations they consist of. For example, and to show the notation, we would like to compare between the actual combined perturbation +S−T and the sum of the simple perturbations (+S)+(−T).

Figure fig.add shows all comparisons. In each case, the actual combined perturbation elicits responses very similar to the sum of responses of the corresponding simple perturbations, both for small and large perturbations (no significant differences in any case). This result, typically associated with linear systems, is relevant to build a conceptual model of the error correction mechanism in a way consistent with the previous section.

**Figure fig.add.**
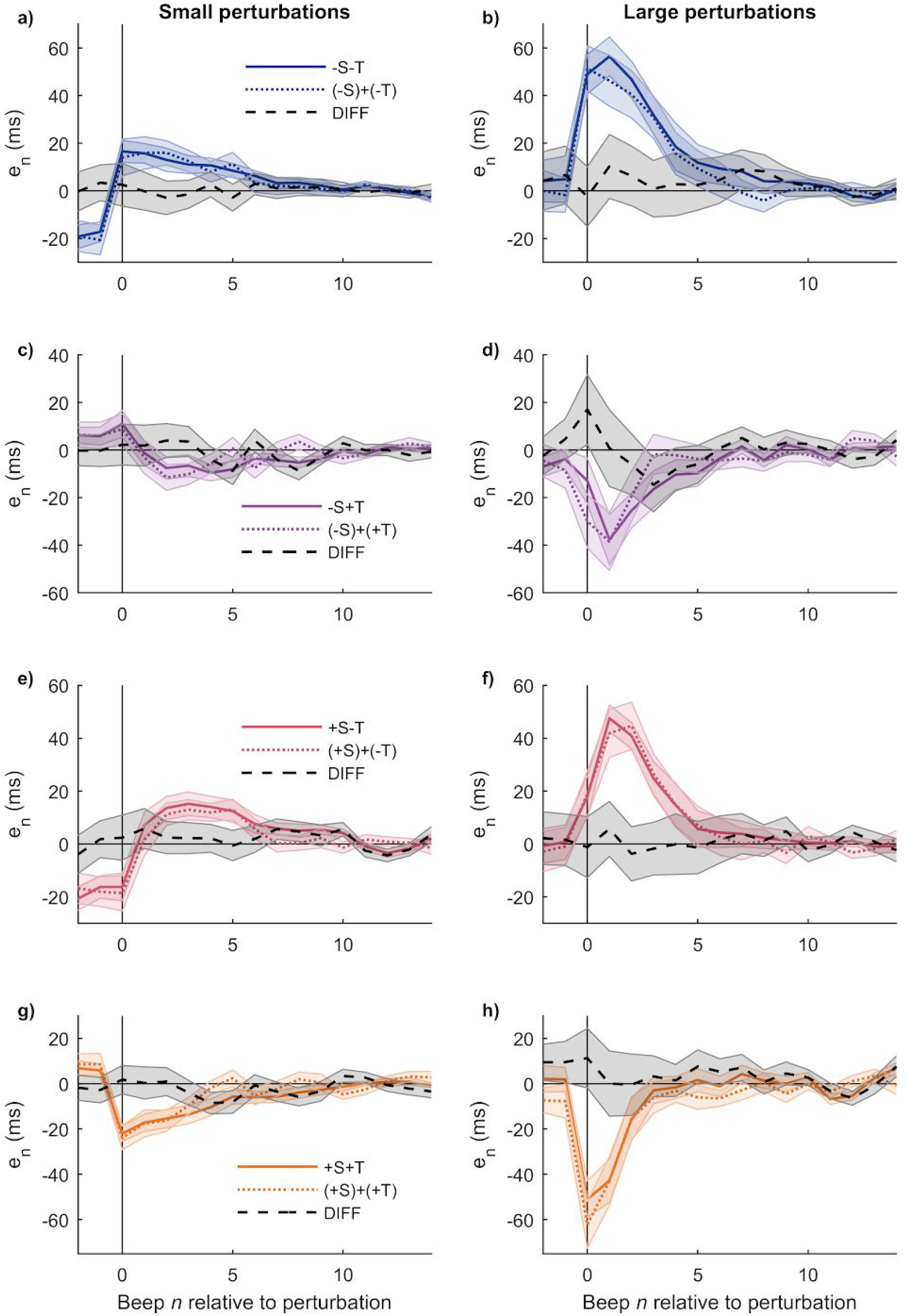
Simple perturbations are additive. Each panel displays the actual combined perturbation, the sum of its individual components, and the difference (DIFF) between the two. (a,b) Combined −S−T and sum (−S)+(−T). (c,d) Combined −S+T and sum (−S)+(+T). (e,f) Combined +S−T and sum (+S)+(−T). (g,h) Combined +S+T and sum (+S)+(+T). Left: small perturbations; Right: large perturbations. The DIFF series are not significantly different from zero in any case. Mean +/− standard error across subjects..

### 2.4. A coarse-grained conceptual model for the error correction mechanism

The results shown in the previous sections allow us to distinguish between various ways of how temporal information is processed in this task and so we will consider a very simple, coarse-grained, conceptual model for the error correction mechanism. The input is the temporal information coming from performance monitoring (time perception) and that could enter the system through different paths depending on the condition--for instance, temporal information from a simple temporal perturbation T and from a simple spatial perturbation S could travel different paths. On the other end, the output is the result of processing such temporal information and drives the motor output. The central part is the time processing itself where hypothetical processes like time comparison, decision making and time prediction and production take place [Grondin2010, Barne2018, Paton2018, IvrySpencer2004, IvrySchlerf2008, Bueti2011, Coull2011], and where the estimation of the appropriate asynchrony correction is made. It should be noted that the parts of this model are not necessarily associated with individual brain regions. The central part of time processing might include sensory components (at its beginning) and motor components (at its end), besides the hypothesized processes for time comparison, decision making, etc. That is, part of the processing itself might take place in sensory and/or motor cortices. We propose the following hypothesis:

1. The processing of temporal information is nonlinear. In particular:

a. the response to symmetric perturbations is not symmetric;
b. the response to a combined perturbation is not equal to the sum of responses of the individual perturbations.
2. The processing of temporal information coming from temporal perturbations and from spatial perturbations is performed by a common mechanism.

Our hypotheses are represented in four versions of the conceptual model as combinations of 1) linear vs. nonlinear processing, and 2) common vs. separate processing. The four combinations are displayed in Figure fig.models. A straightforward result is the observed asymmetric responses, sections 3.1 and 3.2, that validate hypothesis 1.a and thus the linear models (c) and (d) must be discarded. It is worth noting that the symmetric responses to the spatial perturbations S (Figure fig.simplesb) don’t necessarily imply that the processing of asynchronies coming from spatial perturbations is linear and consequently discard hypothesis 1.a--as we noted above, spatial perturbations S in our experiments produced small asynchronies in comparison to the ones produced by the large temporal perturbations T and thus they might be insufficient to elicit typical nonlinear behaviors.

**Figure fig.models.**
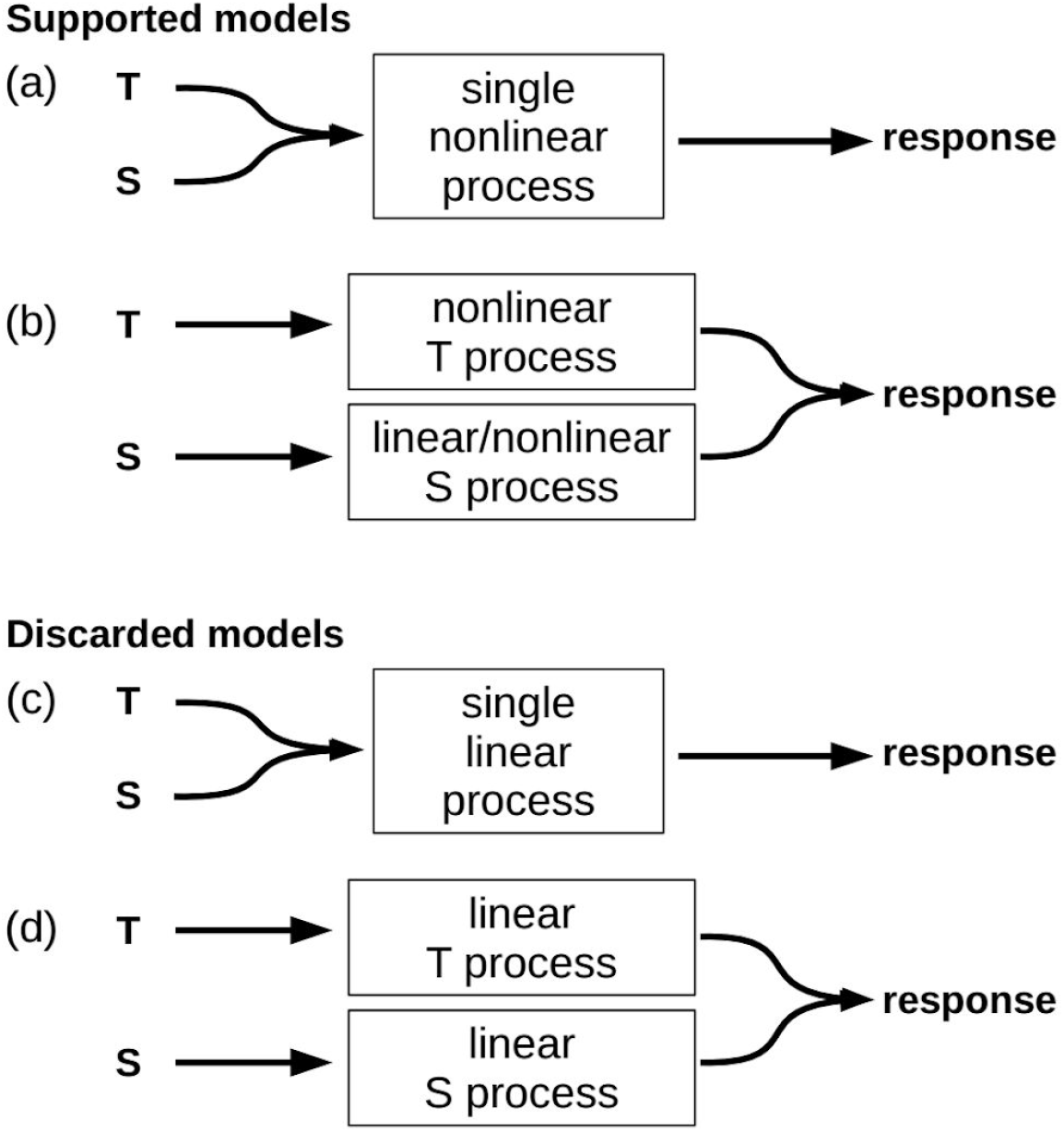
Coarse-grained conceptual models for the processing of temporal information. Models (a) and (c) assume a common mechanism for the processing of temporal information coming from either temporal manipulations T or spatial manipulations S. Models (b) and (d) assume that T and S are processed separately and are added up at the end.

Now consider hypotheses 1.b and 2. An unwarranted conclusion from the additivity results presented in Figure fig.add would be that T perturbations and S perturbations are processed either by a common linear mechanism or by separate nonlinear mechanisms and then added up. However, in front of the discussion of the previous paragraph, if the S perturbations are indeed too small to elicit nonlinear behavior then we would rather expect they might show as additive. That is, the most parsimonious conclusion is that we cannot discard a single nonlinear mechanism and thus we cannot decide whether hypotheses 1.b and 2 are true or false. The two models backed up by our results are then (a) and (b), and to distinguish between them we would have to perform perturbations where the asynchronies produced by S perturbations were comparable in size to those produced by the large T perturbations. However, the remarkably consistent behavior across perturbation sizes, perturbation signs, and perturbation types is noteworthy, suggesting there might be a single underlying mechanism in charge of correcting asynchronies regardless of their origin.

### 2.5. Combined perturbations and exploration of new system states

#### 2.5.1. Experimental phase space

In a previous work, we showed experimental and theoretical evidence suggesting that the correction mechanism underlying resynchronization after a temporal perturbation in paced finger tapping is bidimensional and nonlinear [Bavassi et al 2013]. If one of the variables of such a system is the observable *e*_*n*_, how could we obtain information about the other variable? The embedding technique [Gilmore1998] allows us to get information about the geometric organization of the underlying system’s trajectories by analyzing the time series of a single observable, in this case *e*_*n*_.

To illustrate the procedure, in Figure fig.embedT we undo the “small/large” classification of perturbation sizes and show the averaged time series corresponding to the simple temporal perturbations T of all sizes and signs (+/−15ms, +/−30ms, +/−45ms, +/−50ms; panel A). In addition, we show in panel B the trajectories that result after reconstruction of the experimental phase space by means of an embedding where the first component is ½(*e*_*n*_ + *e*_*n−1*_) and the second one is ½(*e*_*n*_ − *e*_*n−1*_). The perturbation step, n=0 in the time series, corresponds to the initial dot in the trajectory in the embedding, and the origin represents the stationary solution after resynchronization where all asynchronies are zero on average. The trajectories are neatly organized according to size and sign of the perturbation. Responses to larger perturbations begin farther away from the origin; the trajectories for the 45 ms and 50 ms perturbations are very similar as expected by the similitude between the perturbation sizes. The clear geometric organization of the embedded trajectories supports the idea of a common error correction mechanism in charge of resynchronization whose response depends on the size and sign of the perturbation [Bavassi2013].

**Figure fig.embedT.**
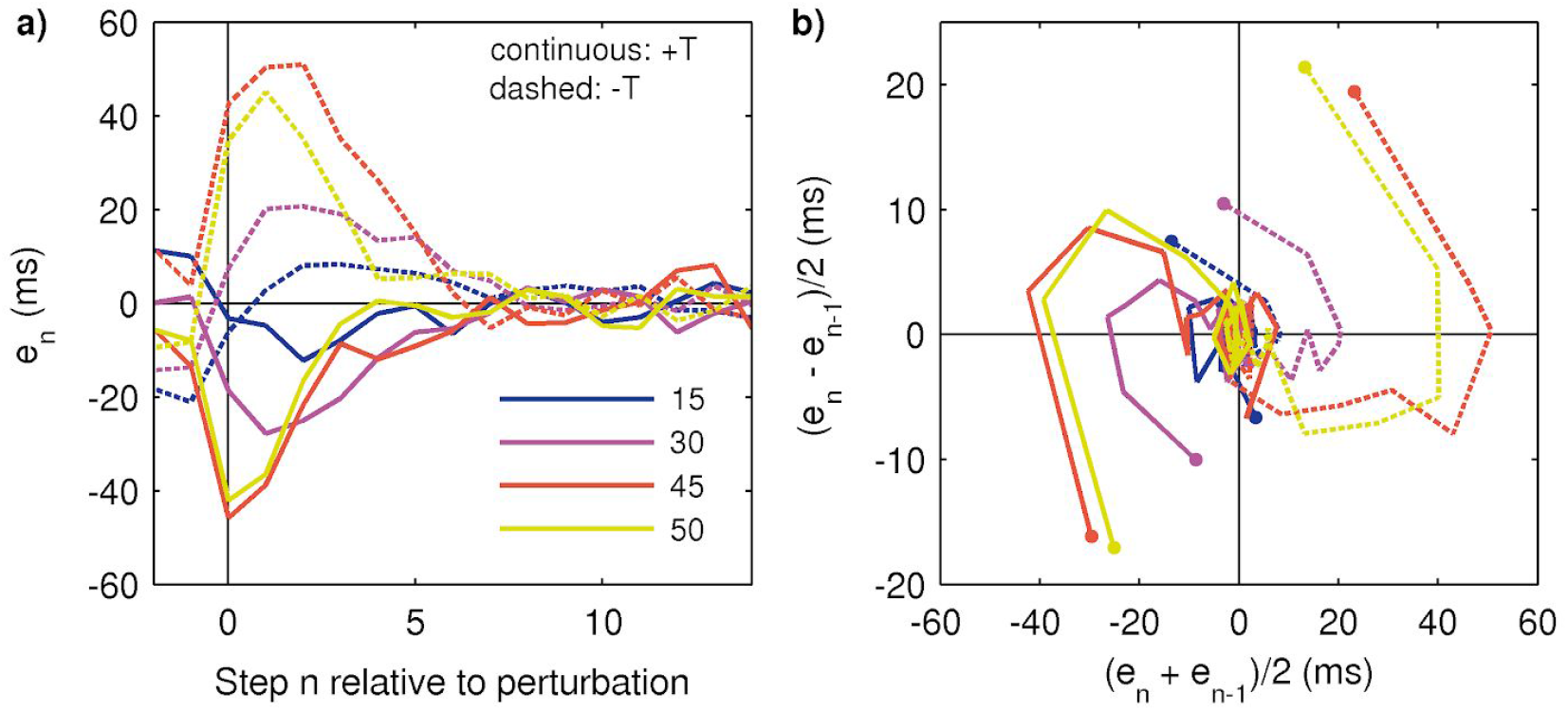
Experimental reconstruction of the phase space by means of an embedding. (a) Averaged time series of *e*_*n*_ in response to temporal perturbations T of all sizes and signs. (b) Embedding of the time series shown in (a). The trajectories are organized in phase space according to the size and sign of perturbation (the smallest perturbations +/−15ms follow the same organization than the larger ones but are visually masked by the post-perturbation variability). The beginning of each trajectory is marked by a dot and corresponds to the perturbation step n=0 in the time series (steps before the perturbation are not shown for visual clarity). All trajectories converge to the origin, representing the post-perturbation baseline. Mean across subjects; the error bands are not shown for visual clarity.

#### 2.5.2. Trajectories of the combined perturbations

We’ve shown that the simple perturbations S produce a novel effect: a change in asynchrony without a change in the stimulus period. On the other hand, the combined opposite perturbations (+S−T and −S+T) are also novel: they produce a change in the period without changing on average the expected asynchrony value (see Methods and Supplementary Figure fig.schematic). For instance, let’s consider a traditional simple temporal perturbation-T representing a tempo step decrease (a change in stimulus period by an amount −DeltaT). It normally produces an increase in asynchrony by an amount −DeltaT at the perturbation step. By combining −T with +S to get a +S−T perturbation, a decrease in stimulus period is produced due to the −T component but with a lesser asynchrony value because of the temporal compensation produced by the +S component. Conversely, the combined analog perturbations (+S+T and −S−T) make the asynchrony value greater (in absolute value) than the one produced by a T perturbation alone.

In Figure fig.embedSTa we compare the time series of the traditional perturbations −T and +T (size 50 ms only) and the corresponding combined opposite and analogous perturbations. All three types of perturbations (traditional, opposite, analogous) produce similar time evolutions except for their initial asynchrony value (n=0). In the phase space, this can be seen as the three trajectories on the same side making similar paths after a quick convergence from differing initial points. That is, the responses to perturbations that have a change in period are similar despite different initial asynchrony values.

**Figure fig.embedST.**
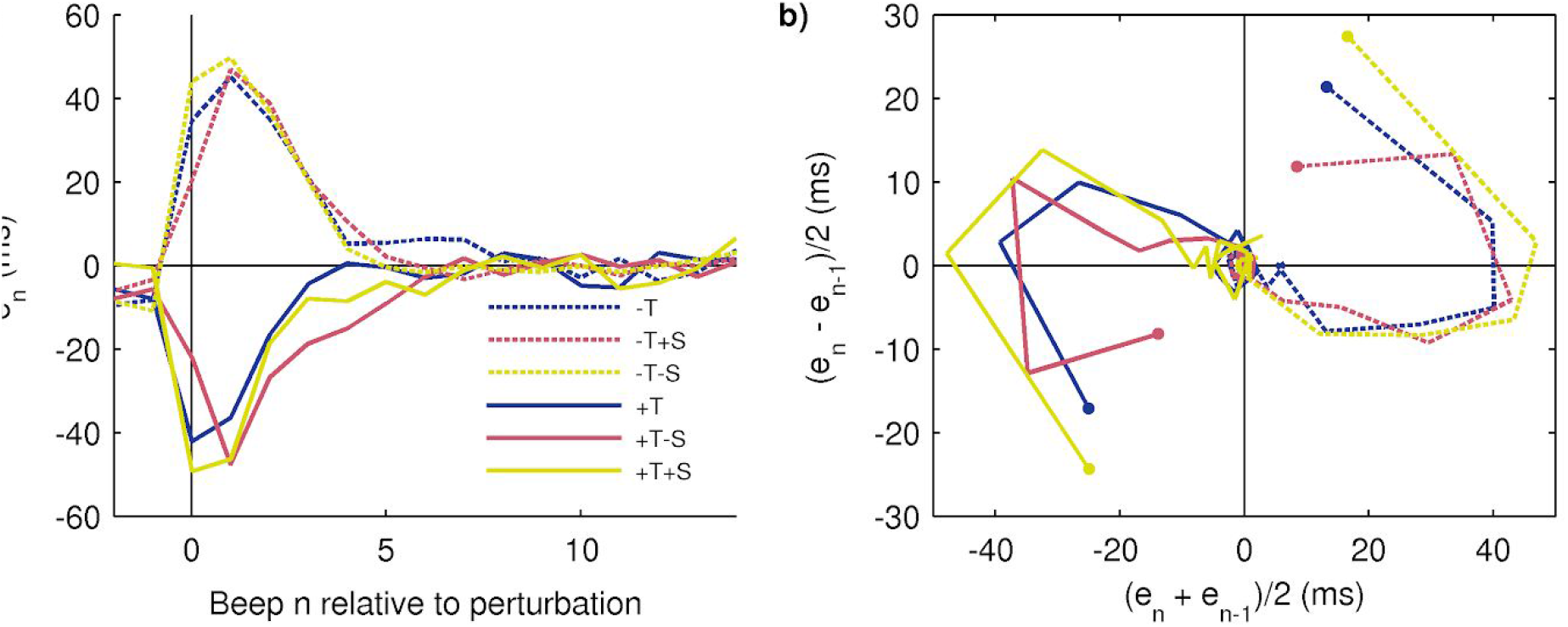
Response to combined perturbations of +/−50 ms and experimental reconstruction of phase space. (a) Averaged time series of traditional simple temporal perturbations (+T and −T), combined opposite perturbations (−S+T and +S−T), and combined analogous perturbations (+S+T and −S−T). (b) Embedding of the time series shown in (A); the dots correspond to the perturbation step (n=0; previous steps are not shown for visual clarity). The paths taken by the opposite and analogous perturbations are very similar to the simple temporal perturbations but with different initial points; the initial points are organized according to the expected asynchrony value (from the origin outwards: opposite-simple-analogous). Mean across subjects; the error bands are not shown for visual clarity.

#### 2.5.3. All perturbations: new system states

In this section we continue analyzing the experimental phase space shown above by including the information from the novel perturbations presented in this work. The combined perturbations, as well as the already discussed simple spatial perturbations S, allow us to decouple the effect of traditional temporal perturbations and access as yet unexplored system states. We will normalize the trajectories from all perturbations so we can make evident their positions in phase space relative to one another.

Figure fig.areasa shows the average of simple temporal perturbations +/−T (averaging across 45 and 50 ms sizes only for clarity, and after normalization and non-dimensionalization, see Methods). Shaded areas represent the regions in phase space (i.e. system states) explored by the traditional tempo step change perturbations. In Figure fig.areasb we include the rest of the perturbations (simple spatial +/−S, combined opposite +/−S-/+T, and combined analogous +/−S+/−T, after normalization and non-dimensionalization). It is evident that the regions in phase space explored by the novel perturbations overlap with the traditional +/−T but only partially, and thus they allow us to probe the system in novel ways and access system states as yet unexplored as a result of decoupling the experimental manipulation of period and asynchrony.

**Figure fig.areas.**
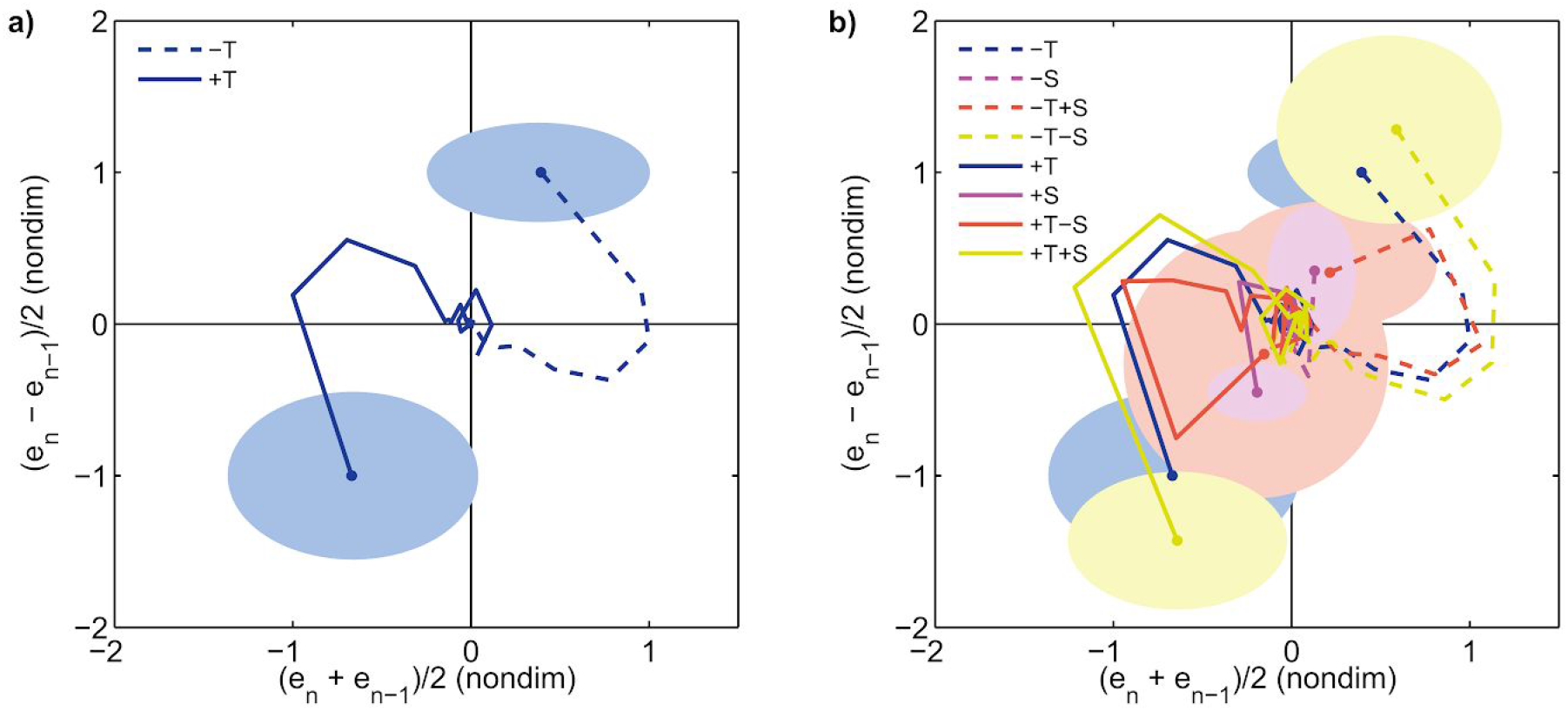
Access to previously unexplored system states. (a) Embedding of the responses to traditional perturbations +/−T in normalized, non-dimensionalized, averaged coordinates (as before, the displayed trajectories start at the perturbation step marked by a dot, corresponding to n=0 in the time series). The shaded areas correspond to the region in phase space where the system is left at the perturbation step (center: mean across subjects; radii: standard deviation across subjects). (b) Embedding of all perturbations. The novel perturbations probe the system in new ways by setting initial conditions (i.e. at the time of perturbation) never tested before; this is evidenced by the areas corresponding to novel perturbations only partially overlapping with the traditional perturbation.

## 3. Conclusions

Experimental work on sensorimotor synchronization and, in particular, both paced and unpaced finger tapping date back to the end of the XIX century [Stevens1886], while the first mathematical models were published 50 years ago, e.g. [Michon1967]. Much work has been done in this area of research to advance our knowledge about the underlying mechanism responsible for achieving average synchrony [Repp2005, Repp2013]. In particular, perturbations to the sequence period are a usual way to probe the system’s inner workings. As we described in this work, these perturbations are in fact confounded parameter and variable manipulations and, up to our knowledge, no published work has addressed this issue.

Our observation applies to any perturbation to the sequence period: local step changes, phase shifts, and event onset shifts, and global changes like accelerando or ritardando, etc. A different, less usual way of perturbing the system is also worth discussing: temporally delayed or advanced auditory feedback from the taps [AscherPrinz1997, Wing1977, PfordDalla2011, MatesAscher2000]. Note that the period of the sequence is not changed in this kind of perturbation. However, it introduces a dissociation between auditory feedback and proprioceptive and tactile feedback from the taps and thus its relationship to the (perturbed) asynchrony value remains to be elucidated.

We showed that an experimental phase space can be reconstructed from the averaged time series via embedding and that all trajectories (corresponding to either simple or combined perturbations of any size and sign) organize in a geometrically remarkable way. This supports the notion that the underlying system in charge of the error correction can be considered as a single mechanism [Bavassi2013], as opposed to many different mechanisms that are turned on or off depending on the magnitude, sign, and type of perturbation, a usual---and frequently implicit---assumption when perturbation data are considered [Thaut1998]. Note that this doesn’t imply that the details of such a mechanism are simple---in fact, we expect that the hypothetical processes within such a mechanism like time perception, comparison, and production interact in complex ways, particularly in this task where processing stimuli and producing timed responses is ongoing and the processes related to step *n* are likely overlapped in time with those from the previous step *n*−1 [Bavassi2017]. Yet, according to our results, it appears that the response of the system is consistent regardless of whether the asynchrony has a purely temporal or spatiotemporal origin, and of the size and sign of the perturbation.

Research about how temporal information is processed in the brain has grown very rapidly in the last decade, particularly research related to the neural underlying mechanisms and involved brain regions [Paton2018]. Many important issues remain open, like whether timing is centralized and dedicated or distributed or intrinsic; whether it reflects properties of individual neurons or is the emergent behavior of neural networks; whether sensory and motor timing rely on the same circuitry or not; and so on. Our results show that traditional perturbations are confounded parameter-variable manipulations, and that a broader set of perturbations (to either the parameter, the variable, or both) leave the system in different points of the phase space. This suggests revisiting the usual assumptions about perturbations and interpreting results with this distinction in mind. In the music domain, for instance, our results open the possibility that different brain regions and processes might be recruited in front of a perturbation to either the asynchrony or the tempo [Zatorre2007], and that period and asynchrony might have different neural representations or correlates. Whether the known associations between functions and regions would split after solving the confounding is yet to be elucidated.

## 4. Methods

### 4.1. Task and subjects

The task was an auditorily paced finger tapping that may, at some random point in the series, present a perturbation. The subject was instructed to keep in synchrony at his/her best by using his/her index finger (holding the wrist in place during the experiment) and keep tapping to resynchronize in case a perturbation appears. Each trial was either isochronous (constant stimulus period and fixed point of contact) or perturbed. Perturbations were any of the following three types: a) simple temporal perturbations “T” (traditional tempo step change perturbations); b) simple spatial perturbations “S” that had a temporal effect (see below); c) combined simultaneous perturbations “ST”. Subjects were volunteers; one subject was not able to complete the task, and three subjects were excluded according to the criteria below. The final number of subjects was N=30 (ages 19-40, mean 29.1, 13 female, 28 right-handed, all subjects used their dominant hand).

### 4.2. Perturbations

We performed three types of perturbations:

a. Simple temporal perturbations +/−T. These perturbations were the traditional tempo step changes where the stimuli period changes once by an amount +/−DeltaT.
b. Simple spatial perturbations +/−S. These novel perturbations consisted in raising (+) or lowering (−) the contact point between finger and sensor, which had the temporal effect of advancing or delaying the time of contact (when the sensor was in its higher or lower position, respectively) and thus made the asynchrony at the perturbed step more negative or more positive, respectively.
c. Combined perturbations ST, where simultaneous (i.e. at the same step n) S and T perturbations take place: +S+T, −S+T, +S−T, −S−T.

In any case the perturbation was unexpected (it occurred at a randomly chosen step).

### 4.3. Setup

The setup consisted of two interconnected microcontrollers (Arduino Mega) communicating via the Wire library. The “master” microcontroller received instructions from the control script (MATLAB, The MathWorks, Inc) and started a trial (it received trial parameters, generated the sequence of auditory stimuli, sent instructions to the slave microcontroller, registered taps, sent trial data back to the control script). The master microcontroller had a custom-designed shield [Bavassi2017] for interfacing and signal conditioning (sum and amplification of audio signals, virtual ground, voltage divider for the force sensor, etc.). Every time a tap was detected, auditory feedback was sent as a brief tone. Stimuli and feedback tones were 50-ms duration square waves of 440 Hz (A4) and 587 Hz (around D5) respectively. Visual feedback from the subject’s hand was avoided by means of a blocking screen. The “slave” microcontroller was in charge of producing the spatial perturbations S whenever the master sent the instruction. The slave microcontroller drove a small platform by means of a servo motor (Savox SC-1258TG Super Speed Titanium Gear Standard Digital) and a rotating arm, displacing the platform upwards or downwards along a linear ball bearing. The platform had a force-sensitive resistor (FSR 406, Interlink Electronics) attached on top of it to detect the subject’s response. Sound was played diotically through Sennheiser HD419 headphones, and subjects adjusted sound volume to a comfortable level.

### 4.4. Experimental design

The combination of temporal perturbations T, spatial perturbations S, and perturbation direction (positive or negative) resulted in 9 experimental conditions: isochronous (no perturbation), 4 simple perturbations (temporal +T and −T and spatial +S and −S), and 4 combined perturbations (+S+T, −S+T, +S−T, −S−T). Each subject participated in a single session with three phases: Introduction, Calibration, Test. Each phase ended when all trials were successfully completed. Each trial consisted of a sequence of 30 auditory stimuli with a period (interstimulus interval) of 500 ms until a perturbation occurred randomly between the 15th and the 20th stimuli. In the temporal perturbations the period of the sequence was either increased (positive, +T) or decreased (negative, −T). In the spatial perturbations the vertical position of the platform was either moved upwards (positive, +S) or downwards (negative, −S) from its resting position by a distance that depended on the subject (see calibration below) and the new position was kept until the end of the trial. In the combined perturbations both T and S manipulations were performed simultaneously. The introduction phase was an exposure to the task and the 9 experimental conditions (one trial per condition, 9 trials total) to allow the participant to familiarize with the experimental task. The calibration phase consisted of 8 trials with simple spatial perturbations +/−S with the most extreme positions of the platform (+/−1.5 cm, 16 trials total), so we can estimate the longest temporal effect of the spatial perturbations for each subject. The test phase consisted of 8 trials from each experimental condition (72 trials total), in three blocks separated by short breaks to prevent the participant from becoming tired. Trials were presented in random order within each phase.

### 4.5. Calibration

The temporal effect of raising or lowering the platform is subject-dependent since every subject moves the finger at its own speed and, thus, a given spatial displacement of the platform is translated to different times for the finger to reach the new contact point. In the calibration phase we estimated for each subject the average time error produced by the unexpected movement of the platform from its resting position to its two most extreme positions (highest 1.5 cm, lowest −1.5 cm).

### 4.6. Experiments

We performed two experiments:

#### Experiment 1

Matched temporal and spatial perturbations (i.e., the temporal effect of the spatial perturbation is equal to the assigned temporal perturbation size). Subjects were assigned to either one of the following groups, depending on the recorded average time error during the calibration phase:

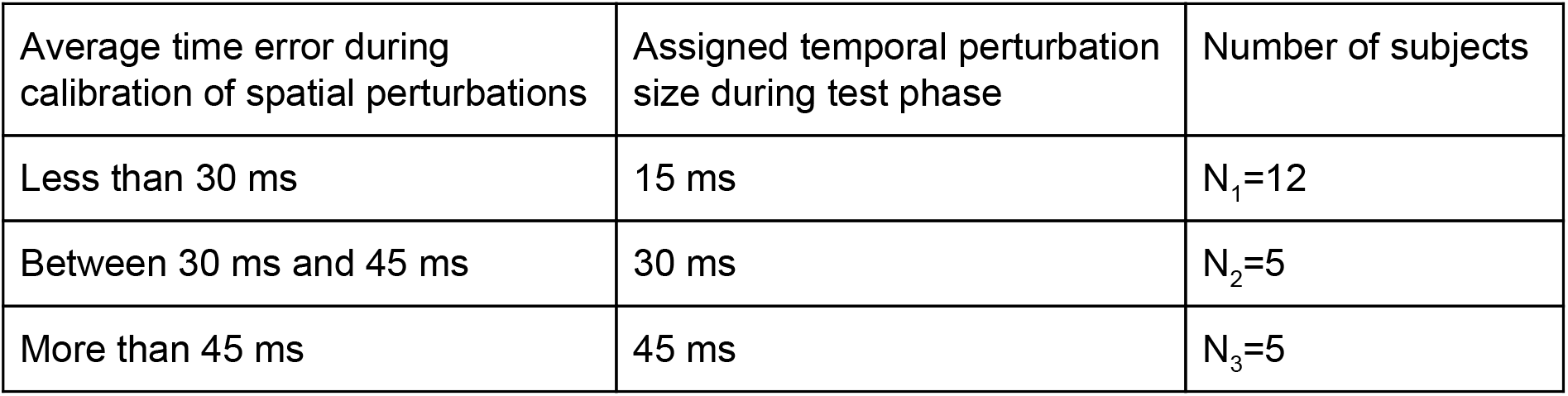

| Average time error during calibration of spatial perturbations | Assigned temporal perturbation size during test phase | Number of subjects |
| --- | --- | --- |
| Less than 30 ms | 15 ms | $N_1=12$ |
| Between 30 ms and 45 ms | 30 ms | $N_2=5$ |
| More than 45 ms | 45 ms | $N_3=5$ |

#### Experiment 2

Unmatched temporal and spatial perturbations. Subjects with a large temporal effect in spatial perturbations were very infrequent, so in order to test larger temporal perturbation magnitudes we created a fourth category with +/−50-ms temporal perturbations and +/−15-ms spatial perturbations. Number of subjects: N_4_=8.

### 4.7. Exclusion criteria

A trial was correct if the subject began tapping before the 6th stimulus and if the subject didn’t miss any response from there on. Each incorrect trial was repeated until completed successfully. Asynchrony outliers: after finishing the experiment, any trial with very large asynchronies (greater than 145 ms in absolute value, with respect to the pre-perturbation baseline) was discarded. Standard deviation outliers: the distribution of the standard deviation of pre-perturbation asynchronies for each subject was computed, and any trial with a standard deviation greater than 1.5 times the IQR was discarded. After discarding asynchrony and deviation outliers, subjects with less than 4 correct trials for every condition were removed from further analysis (3 subjects, 3 different conditions).

### 4.8. Data pre-processing

In order to average across trials and subjects, we aligned all trials at the perturbation step (renamed as n=0) and defined the analysis range from n=−10 to n=14 such that within that range all subjects have all responses. We define the trial pre-perturbation baseline as the average of all asynchronies between n=−7 and n=−1, so we can avoid adaptation effects at the beginning of the trial. The trial post-perturbation baseline is the average asynchrony between n=9 and n=14.

### 4.9. Classification of perturbations

When specifically considering perturbations we leave out the isochronous condition, and thus there are 4 perturbation sizes (15, 30, 45, 50 ms) and 8 conditions (+T, −T, +S, −S, +S+T, −S+T, +S−T, +S+T).

#### According to their size

In order to increase the statistical power, a posteriori we grouped the data in two categories according to the perturbation size: small and large perturbations. Small perturbations include 15-ms and 30-ms perturbation sizes (N=N_1_+N_2_=17 subjects), while large perturbations include 45-ms and 50-ms sizes (N=N_3_+N_4_=13 subjects).

#### According to their type

According to their origin and expected effect, perturbations can be classified as:

Simple: temporal +/−T and spatial +/−S.

Combined: simultaneously changing the period and raising/lowering the point of contact. If the temporal effect of both components is in the same direction (i.e., the asynchrony at the perturbation step either increases or decreases) the perturbation is Analogous (−S−T and +S+T); if the effect goes in opposite directions (for instance one component increases the asynchrony while the other decreases it) the perturbation is Opposite (−S+T and +S−T). See Supplementary Figure fig.schematic.

### 4.10. Averaging

Time series in Figures fig.todas through fig.embedST are grand averages across subjects. Each subject’s average is the mean of all his/her included trials from the corresponding condition.

### 4.11. Hypothesis testing

#### Asymmetry

In order to quantify the degree of asymmetry between responses to symmetric perturbations we define the measure ASYM as the sum of the two corresponding time series (i.e. the algebraic difference between one and the opposite of the other). In this way we can compare time series that have opposite signs by definition (because they result from perturbations with opposite signs) and, at the same time, any series is allowed to change sign in the middle without affecting the analysis (it would be artificially rectified if we otherwise used its absolute value). ASYM values close to zero indicate that the compared time series are mostly symmetric; large positive or negative values indicate asymmetry.

#### Additivity

In order to quantify the additivity between responses to simple perturbations we define the measure DIFF as the difference between the time series of the experimental combined perturbation and the time series obtained by summing the two corresponding individual simple perturbations. For instance, DIFF is the difference between the experimental combined perturbation +S+T and the algebraic sum of the experimental simple perturbations +S and +T. DIFF values close to zero indicate that the combined compared time series are similar; large positive or negative values indicate that the time series are different.

#### False Discovery Rate (FDR) correction

In order to test the statistical significance of ASYM and DIFF we computed p-values for each step between n=1 and n=5 (both included) and then pass the five p-values to the FDR algorithm [BenjaminiHochberg1995] with alpha=0.05. In order to compute the p-values we generated null distributions of ASYM time series and DIFF time series. We illustrate the procedure with an example. To generate the null distribution of ASYM for the small +T/−T comparison (Figure fig.simplesa) we first pooled all trials from all subjects from both conditions +T and −T; then randomly pick 8 trials, averaged them and assigned them to the “surrogate +T” time series of surrogate subject 1; similarly with other 8 trials to the “surrogate −T” of same subject; these surrogate +T and surrogate −T average series were summed to get the surrogate ASYM for this surrogate subject; we then repeated the steps to generate 17 and 13 surrogate subjects for small and large perturbations respectively in order to match the actual number of subjects; the average across surrogate subjects makes one surrogate grand average ASYM time series, and finally repeated the whole procedure 1000 times to get the null distribution of ASYM time series, i.e. the null distribution of ASYM for each step n. Since we are interested in the transient part of the resynchronization, we restricted the FDR analysis to the range n=1 to n=5, both included (n=0 is excluded because it is the perturbation step and the asynchrony there is a forced error; steps n>5 are too close to the post-perturbation baseline). We then compared the true ASYM value at each step between n=1 and n=5 to the null distribution of ASYM at the same step and compute a p-value as the proportion of null ASYM values above the true value or below its opposite (two-tailed). We applied the same procedure to all comparisons between times series in this work.

### 4.12. Phase space reconstruction (embedding)

#### *Regular trajectories* (*Figures* fig.embedT *and* fig.embedST)

Given a univariate time series *e*_*n*_, there are several possible alternatives to implement time series embedding for phase space reconstruction [Gilmore1998], like the usual time-delay embedding (*e*_*n*_; *e*_*n*−1_). We chose the following: the mean between consecutive values (*e*_*n*_+*e*_*n*−1_)/2 for the first variable and the semi-difference (*e*_*n*_−*e*_*n*−1_)/2 for the second variable. Our choice was based on visual clarity and greater trajectory separation (we also tested other embeddings like the above-mentioned time-delay embedding with qualitatively similar results). All reconstructed trajectories in this work were computed directly from the corresponding grand average time series.

#### *Normalized trajectories* (*Figure* fig.areas)

Normalized trajectories were computed by first taking the regular trajectories from 45 and 50 ms perturbation sizes and averaging between them (separately for positive and negative perturbations), then taking each of these and dividing it by its corresponding absolute maximum value (separately for the first and second embedding components). Then we kept the normalized +/−T trajectories as standards and multiplied every trajectory (+/−S, +/−S+/−T, +/−S−/+T) by a factor equal to the ratio between its absolute maximum value and the absolute maximum value of the T trajectories.

#### *Areas* (*Figure* fig.areas)

Areas in Figure fig.areas are a rough measure of the region in phase space where the system is left when the perturbation arrives (step n=0), in units of the normalized trajectories. Areas are ellipses with a center and two radii. The center (which roughly corresponds to the first point of the corresponding normalized trajectory) is the average across subjects of the point in trajectory corresponding to n=0, after pooling 45 and 50 ms perturbation sizes (separately for positive and negative perturbations). The radii are the standard deviation in each component.

### 4.13. Color blind-friendly plots

We used the ametrine colormap [GeissbuehlerLasser2013].

## Acknowledgements

This work was supported by Universidad Nacional de Quilmes (Argentina), CONICET (Argentina), and The Pew Charitable Trusts (USA). We thank Ignacio Spiousas for detailed comments and suggestions on the manuscript.

## Author Contributions

SLL and RL designed the experiments. RL conceived of the approach. SLL wrote the code, built the experimental setup and performed the experiments. SLL and RL analyzed the data. SLL and RL wrote the paper.

## Additional Information

The authors declare no competing interests.

## Supporting Data and Files

The datasets generated and analysed during the current study and the code to plot all figures will be available in the Sensorimotor Dynamics Lab’s website www.ldsm.web.unq.edu.ar/independent2019.

## Supplementary Information

**Figure fig.schematic.**
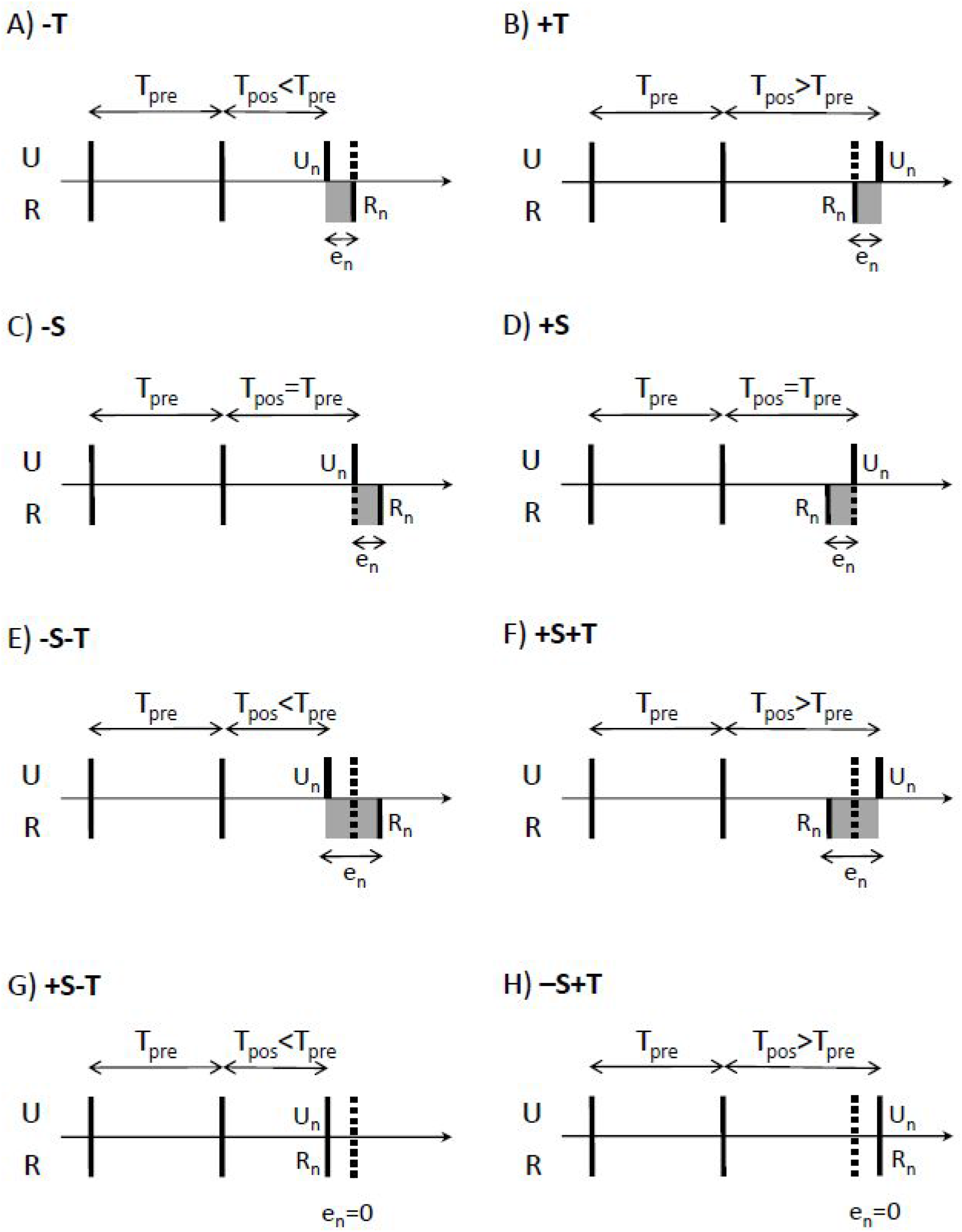
Schematic of perturbations. U = stimuli sequence. R = responses (considering perfect average synchrony for simplicity). Vertical continuous lines: actual occurrences of stimuli and responses. Vertical dashed lines: expected occurrences of stimuli and responses if no perturbation occurs. A vertical dashed line at the level of stimuli means a temporal perturbation occurred. A vertical dashed line at the level of responses means a spatial perturbation occurred. The shaded area indicates the expected average asynchrony at the perturbation step.

**Figure fig.setup.**
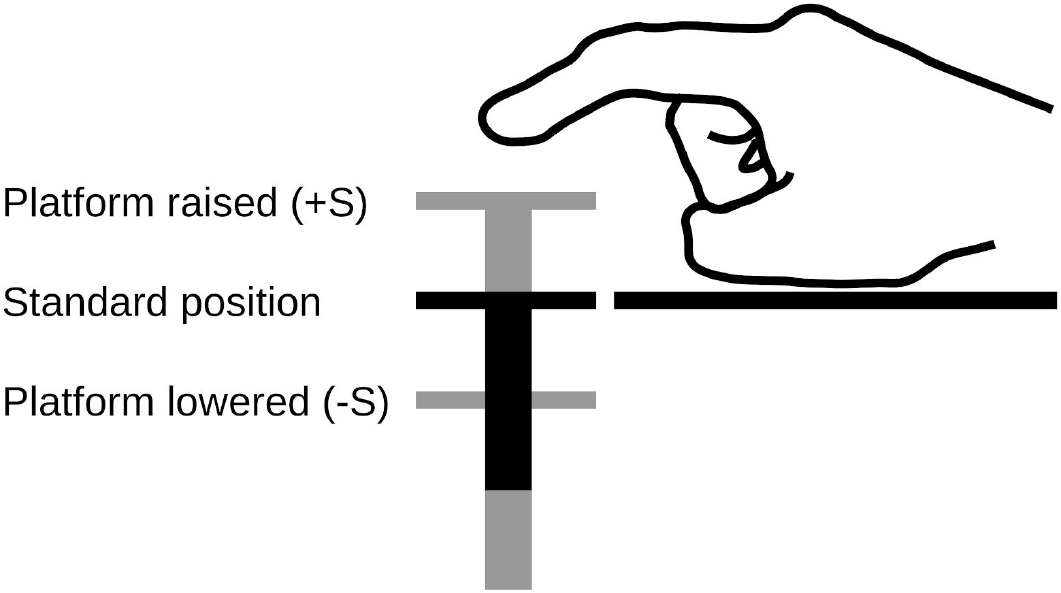
Schematic of setup. The setup consists in a device of three levels. The first level is required to permit the platform to descend. The second level has the servo motor and the spindles of the platform where the sensor force is placed. Here is where the platform descends to force positive asynchronies. In the third level, the subject lays the hand for the tapping task. During the experiment, the index finger is splinting to constraint the articulations involved in the task and the device is covered.

## References

Lewis T. Stevens. On The Time-Sense. Mind 11, pp. 393–404 (1886)

J. A. Michon and N. J. L. Van Der Valk. A DYNAMIC MODEL OF TIMING BEHAVIOR. Acta Psychologica 27 (1967) 204–212.

Aschersleben & Prinz 2997, Delayed Auditory Feedback in Synchronization

Wing 1977, Perturbations of auditory feedback delay and the timing of movement

Pfordresher & Dalla Bella 2011, Delayed Auditory Feedback and Movement

Mates & Aschersleben 2000, Sensorimotor synchronization: the impact of temporally displaced auditory feedback

Robert Gilmore. Topological analysis of chaotic dynamical systems. Rev. Mod. Phys. 70, 1455–1529 (1998).

Diego A. Golombek, Leandro P. Casiraghi, Patricia V. Agostino, Natalia Paladino, José M. Duhart, Santiago A. Plano, Juan J. Chiesa. The times they’re a-changing: Effects of circadian desynchronization on physiology and disease. DOI: 10.1016/j.jphysparis.2013.03.007

Santiago A. Plano, Diego A. Golombek and Juan J. Chiesa. Circadian entrainment to light–dark cycles involves extracellular nitric oxide communication within the suprachiasmatic nuclei. DOI: 10.1111/j.1460-9568.2010.07120.x

Kim, J. K., Forger, D. B., Marconi, M., Wood, D., Doran, A., Wager, T., … Walton, K. M. (2013). Modeling and Validating Chronic Pharmacological Manipulation of Circadian Rhythms. CPT: Pharmacometrics & Systems Pharmacology, 2(7), e57–. http://doi.org/10.1038/psp.2013.34

Bruno H. Repp, Peter E. Keller, Nori Jacoby (2012). Quantifying phase correction in sensorimotor synchronization: Empirical comparison of three paradigms. Acta Psychologica 139, 281–290

Repp, B.H. & Su, YH. Sensorimotor synchronization: A review of recent research (2006–2012) Psychon Bull Rev (2013) 20: 403. https://doi.org/10.3758/s13423-012-0371-2

Aniruddh D. Patel, John R. Iversen, Micah R. Bregman, Irena Schulz. Experimental Evidence for Synchronization to a Musical Beat in a Nonhuman Animal. Current Biology, Volume 19, Issue 10, 2009, Pages 827–830, ISSN 0960-9822, https://doi.org/10.1016/j.cub.2009.03.038.

Adena Schachner, Timothy F. Brady, Irene M. Pepperberg, Marc D. Hauser. Spontaneous Motor Entrainment to Music in Multiple Vocal Mimicking Species. Current Biology, Volume 19, Issue 10, 2009, Pages 831–836, ISSN 0960-9822, https://doi.org/10.1016/j.cub.2009.03.061.

Yanqing Chen, Mingzhou Ding, and J. A. Scott Kelso Long Memory Processes (1/f^alpha Type) in Human Coordination Phys. Rev. Lett. 79, 4501 (1997)

Rodrigo Laje, Patricia V. Agostino and Diego A. Golombek The Times of Our Lives: Interaction Among Different Biological Periodicities Front. Integr. Neurosci., 13 March 2018 | https://doi.org/10.3389/fnint.2018.00010

Diego A. Golombek and Ruth E. Rosenstein. Physiology of Circadian Entrainment. Physiological Reviews 90:3, 1063–1102 (2010)

Michael H. Thaut, Robert A. Miller, Leopold M. Schauer. Multiple synchronization strategies in rhythmic sensorimotor tasks: phase vs period correction. Biol. Cybern. 79, 241–250 (1998)

Loehr, J. D., Large, E. W., & Palmer, C. Temporal coordination and adaptation to rate change in music performance. Journal of Experimental Psychology: Human Perception and Performance, 37(4), 1292–1309 (2011). http://dx.doi.org/10.1037/a0023102

Dean V. Buonomano and Rodrigo Laje. Population clocks: motor timing with neural dynamics. Trends in Cognitive Sciences, Vol. 14, No. 12, 520–527 (2010). doi:10.1016/j.tics.2010.09.002

Grondin, S. Timing and time perception: A review of recent behavioral and neuroscience findings and theoretical directions. Attention, Perception, & Psychophysics 72: 561 (2010). https://doi.org/10.3758/APP.72.3.561

Louise C. Barne, João R. Sato, Raphael Y. de Camargo, Peter M.E. Claessens, Marcelo S. Caetano, and André M. Cravo. A common representation of time across visual and auditory modalities. Neuropsychologia 119, 223–232 (2018).

Joseph J. Paton and Dean V. Buonomano. The Neural Basis of Timing: Distributed Mechanisms for Diverse Functions. Neuron Volume 98, Issue 4, Pages 687–705 (2018).

Richard B Ivry and Rebecca MC Spencer. The neural representation of time. Current Opinion in Neurobiology 14:225–232 (2004).

Richard B. Ivry and John E. Schlerf. Dedicated and intrinsic models of time perception. Trends in Cognitive Sciences, Volume 12, Issue 7, Pages 273–280 (2008).

Domenica Bueti. The sensory representation of time. Front. Integr. Neurosci., 08 (2011) https://doi.org/10.3389/fnint.2011.00034

Jennifer T. Coull. “Discrete Neuroanatomical Substrates for Generating and Updating Temporal Expectations”, in Space, Time and Number in the Brain (Academic Press 2011), Pages 87–101. Editors: Stanislas Dehaene and Elizabeth M. Brannon. https://doi.org/10.1016/B978-0-12-385948-8.00007-4

M. Luz Bavassi, Enzo Tagliazucchi, Rodrigo Laje. Small perturbations in a finger-tapping task reveal inherent nonlinearities of the underlying error correction mechanism. Human Movement Science, Volume 32, Issue 1, Pages 21–47 (2013). https://doi.org/10.1016/j.humov.2012.06.002

Bavassi, L., Kamienkowski, J.E., Sigman, M., Laje, R. Sensorimotor synchronization: neurophysiological markers of the asynchrony in a finger-tapping task. Psychological Research 81: 143 (2017). https://doi.org/10.1007/s00426-015-0721-6

Matthias Geissbuehler and Theo Lasser, “How to display data by color schemes compatible with red-green color perception deficiencies,” Opt. Express 21, 9862–9874 (2013).

Benjamini, Y., & Hochberg, Y. (1995). Controlling the false discovery rate: a practical and powerful approach to multiple testing. Journal of the Royal statistical society: series B (Methodological), 57(1), 289–300.

Summer K. Rankin, Edward W. Large, Philip W. Fink. Fractal Tempo Fluctuation and Pulse Prediction. Music Perception: An Interdisciplinary Journal, Vol. 26 No. 5, June 2009; (pp. 401–413) DOI: 10.1525/mp.2009.26.5.401

Madison, Guy, Gouyon, Fabien, Ullén, Fredrik, Hörnström, Kalle. Modeling the tendency for music to induce movement in humans: First correlations with low-level audio descriptors across music genres. Journal of Experimental Psychology: Human Perception and Performance, Vol 37(5), Oct 2011, 1578–1594.

Geoff Luck, Petri Toiviainen. Ensemble Musicians’ Synchronization With Conductors’ Gestures: An Automated Feature-Extraction Analysis. Music Perception: An Interdisciplinary Journal, Vol. 24 No. 2, December 2006; (pp. 189–200) DOI: 10.1525/mp.2006.24.2.189

Pfordresher, P. Q. & Palmer, C. Effects of hearing the past, present, or future during music performance. Percept. Psychophys. 68, 362–376 (2006).

Large, E. W. & Palmer, C. Perceiving temporal regularity in music. Cogn. Sci. 26, 1–37 (2002).

Large, E.W., Fink, P. & Kelso, S.J. Tracking simple and complex sequences. Psychological Research (2002) 66: 3. https://doi.org/10.1007/s004260100069

Levitin, D. J., Grahn, J. A., & London, J. (2018). The Psychology of Music: Rhythm and Movement. Annual Review of Psychology, 69(1), 51–75. doi:10.1146/annurev-psych-122216-011740

Gamma-Band Responses to Perturbed Auditory Sequences: Evidence for Synchronization of Perceptual Processes. Theodore P. Zanto, Edward W. Large, Armin Fuchs, J. A. Scott Kelso. Music Perception: An Interdisciplinary Journal, Vol. 22 No. 3, Spring 2005; (pp. 531–547) DOI: 10.1525/mp.2005.22.3.531

Robert J. Zatorre, Joyce L. Chen & Virginia B. Penhune. When the brain plays music: auditory–motor interactions in music perception and production. Nature Reviews Neuroscience volume 8, pages 547–558 (2007).

Niek R. van Ulzen, Claudine J.C.Lamoth, Andreas Daffertshofer, Gün R.Semin, Peter J. Beek. Stability and variability of acoustically specified coordination patterns while walking side-by-side on a treadmill: Does the seagull effect hold? Neuroscience Letters, Volume 474, Issue 2, 26 April 2010, Pages 79–83 https://doi.org/10.1016/j.neulet.2010.03.008

Roerdink, M., Lamoth, C. J. C., van Kordelaar, J., Elich, P., Konijnenbelt, M., Kwakkel, G., & Beek, P. J. (2009). Rhythm Perturbations in Acoustically Paced Treadmill Walking After Stroke. Neurorehabilitation and Neural Repair, 23(7), 668–678. doi:10.1177/1545968309332879

Pelton, T. A., Johannsen, L., Chen, H., & Wing, A. M. (2010). Hemiparetic Stepping to the Beat: Asymmetric Response to Metronome Phase Shift During Treadmill Gait. Neurorehabilitation and Neural Repair, 24(5), 428–434. https://doi.org/10.1177/1545968309353608

Julia T Choi & Amy J Bastian. Adaptation reveals independent control networks for human walking. Nature Neuroscience volume 10, pages 1055–1062 (2007)

Joseph J. Paton, Dean V. Buonomano. The Neural Basis of Timing: Distributed Mechanisms for Diverse Functions. Neuron, Volume 98, Issue 4, 16 May 2018, Pages 687–705. https://doi.org/10.1016/j.neuron.2018.03.045

M.C. (Marieke) van der Steen, Nori Jacoby, Merle T. Fairhurst, Peter E. Keller. Sensorimotor synchronization with tempo-changing auditory sequences: Modeling temporal adaptation and anticipation. Brain Research, Volume 1626, 11 November 2015, Pages 66–87 https://doi.org/10.1016/j.brainres.2015.01.053

